# Fractal analysis of muscle activity patterns during locomotion: pitfalls and how to avoid them

**DOI:** 10.1101/2020.04.24.059618

**Authors:** Alessandro Santuz, Turgay Akay

## Abstract

Time-dependent physiological data, such as electromyogram (EMG) recordings from multiple muscles, is often difficult to interpret objectively. Here, we used EMG data gathered during mouse locomotion to investigate the effects of calculation parameters and data quality on two metrics for fractal analysis: the Higuchi’s fractal dimension (HFD) and the Hurst exponent (H). A curve is fractal if it repeats itself at every scale or, in other words, if its shape remains unchanged when zooming in the curve at every zoom level. Many linear and nonlinear analysis methods are available, each of them aiming at the explanation of different data features. In recent years, fractal analysis has become a powerful nonlinear tool to extract information from physiological data not visible to the naked eye. It can present, however, some dangerous pitfalls that can lead to misleading interpretations. To calculate the HFD and the H, we have extracted muscle synergies from normal and mechanically perturbed treadmill locomotion from the hindlimb of adult mice. Then, we used one set per condition (normal and perturbed walking) of the obtained time-dependent coefficients to create surrogate data with different fluctuations over the original mean signal. Our analysis shows that HFD and H are exceptionally sensitive to the presence or absence of perturbations to locomotion. However, both metrics suffer from variations in their value depending on the parameters used for calculations and the presence of quasi-periodic elements in the time series. We discuss those issues giving some simple suggestions to reduce the chance of misinterpreting the outcomes.

**New & Noteworthy:** Despite the lack of consensus on how to perform fractal analysis of physiological time series, many studies rely on this technique. Here, we shed light on the potential pitfalls of using the Higuchi’s fractal dimension and the Hurst exponent. We expose and suggest how to solve the drawbacks of such methods when applied to data from normal and perturbed locomotion by combining *in vivo* recordings and computational approaches.

## Introduction

The collection of physiological data over different time spans is crucial for neuroscience research (Sherrington 1906). However, after collecting the data there always remains the issue of finding convenient and informative ways of analyzing it. For instance, electromyogram (EMG) recordings from multiple muscles are widely used for their important role in the study of locomotion (Bizzi et al. 2008). Despite their intuitive meaning (a muscle is either active or inactive and thus either does or does not produce a recordable signal), EMG recordings contain complex features that might not always be detectable and/or quantifiable by traditional analysis methods. To uncover these local or global features of the signal, many linear and nonlinear approaches exist (Kantz and Schreiber 2004; Ting et al. 2015). Here, we used linear decomposition to extract time-dependent motor primitives from EMG data (Santuz et al. 2019). Then, we selected two nonlinear metrics derived from fractal analysis and endeavored to answer two main questions: 1) are those metrics able to detect differences in EMG-derived signal (i.e. motor primitives) when external perturbations are added to locomotion? And 2) what are the potential pitfalls – and ways to avoid them – of using those metrics?

Nature exhibits similar patterns at different scales. A snowflake, when looked through a magnifying glass, shows shapes comparable to those visible to the naked eye. And so do the branches of a tree, the coastline of an island, the course of a river, and many other natural or artificial entities (Mandelbrot 1983). Using mathematics, it is possible to build some geometric constructs that have the property of repeating themselves at every level of zoom, again and again. In other words, they are self-similar (Mandelbrot 1983). Such constructs have been called “fractals” by the mathematician Benoit B. Mandelbrot (Mandelbrot 1983). A celebrated example of fractal, the famous Koch curve (Von Koch 1904) was discovered by another mathematician, Helge von Koch. As shown in the detail of Fig. 1A, the Koch curve repeats itself at every level of zoom: a demonstration of self-similarity. Mandelbrot, following the empirical work of yet another mathematician, Lewis Fry Richardson, asked himself whether it would be possible to quantify the “degree of complication” of such curves (Mandelbrot 1967; Richardson 1961). If in Euclidean geometry a line has dimension 1 and a plane has dimension 2, why not assigning a “fractional” (or fractal) dimension to a curve that is neither a straight line nor, say, a filled rectangle (Mandelbrot 1967)? Since the seminal work of Mandelbrot, the term “fractal analysis” has been used to denote a wide range of approaches, some of them pre-existing, that deal with self-similar entities (Mandelbrot 1983).

**Fig. 1.**
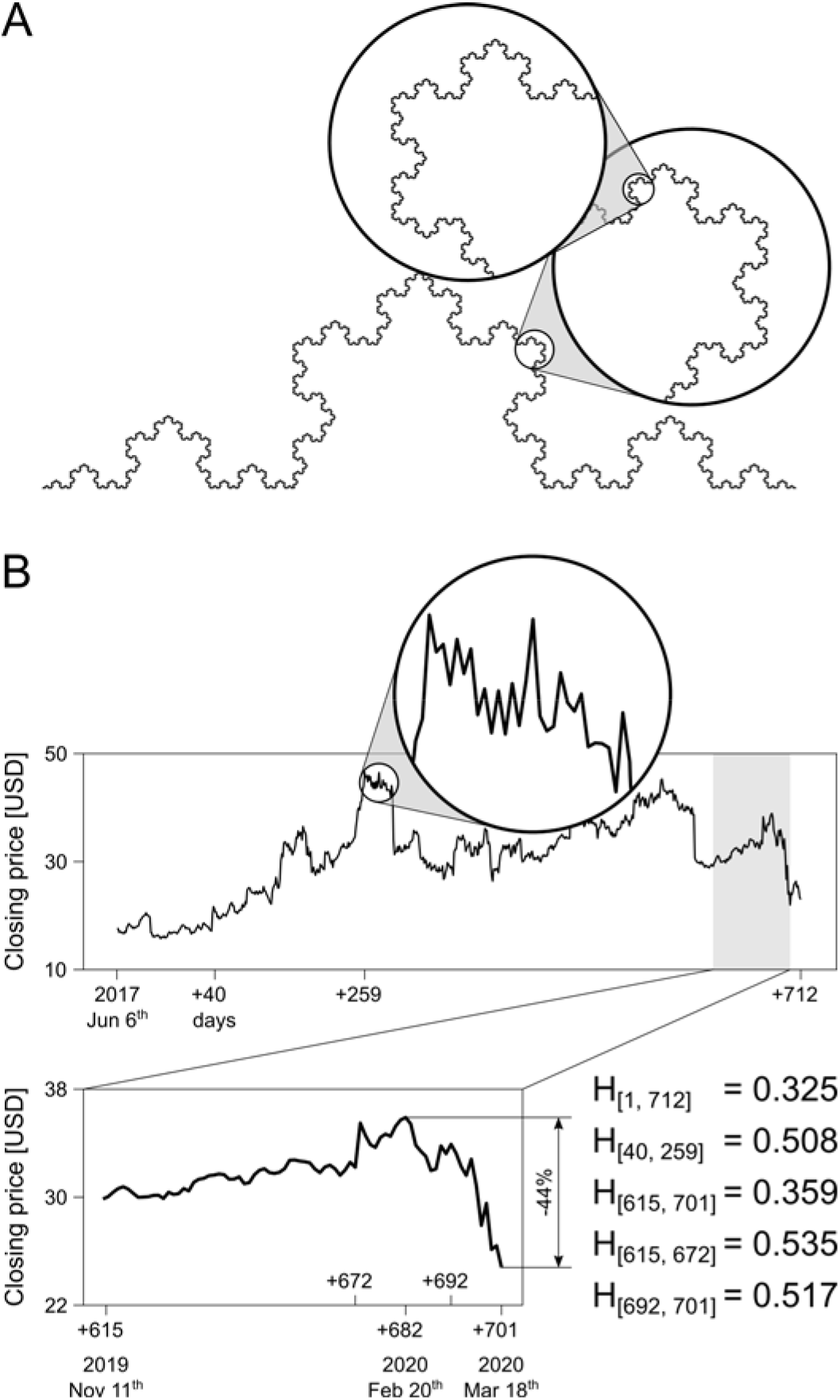
A. The Koch curve is a self-similar curve and it reapeats itself at all scales, as shown in the circular detail. B. Daily market closing price for Twitter stocks (NASDAQ Composite 2020). This time series is not strictly self-similar, but it is self-affine, as shown in the circular detail. The Hurst exponent (H) is calculated for several intervals indicated as days in the subscripts, between brackets. For the calculation of H, the smallest window was set to half the length of the considered interval. 0.5 < H < 1 indicates persistence (negative or positive trend, e.g. that of the inreasing prices between day 615 and 672), while 0 < H < 0.5 indicates anti-persistence (no trend or oscillation around a mean value, e.g. when considering the whole time series from day 1 to 712 or in the case of EMG recordings from locomotion).

Fractal analysis has been used, amongst other disciplines, in hydrology (Hurst 1951), geography (Mandelbrot 1967), genetics (Peng et al. 1994), neuroscience (Lutzenberger et al. 1992), finance (Mandelbrot 1997), physiology and medicine (Losa et al. 2005), biomechanics (Scafetta et al. 2009), and seismography (Flores-Marquez et al. 2012). For instance, it can be used to predict stock market shifts and an example is given in Fig. 1B. Here it is shown the daily stock closing price (NASDAQ Composite 2020) of Twitter, a popular social media amongst scientists (Cheplygina et al. 2020). Due to the SARS-CoV-2 and related COVID-19 outbreak (WHO 2020), after around 60 working days of positive trend, the stock price started to oscillate only to fall drastically by 44% in just 19 working days. The Hurst exponent (H, a metric for fractal analysis, see details in the methods and caption of Fig. 1) values reported in Fig. 1B, show the increasing and decreasing trends before and after the COVID-19 crisis with 0.5 < H < 1. However, when the time series is analyzed as a whole and has many oscillations as those we might find in EMG recordings from locomotion (see e.g. Fig. 2A-B), H values do not display any trend and 0 < H < 0.5. This is an indication that different fractal dimensions can be attributed to the same curve, depending on the scale and subset considered. The discrepancy can be explained by the fact that data coming from real-world scenarios cannot always be classified as self-similar (Mandelbrot 1985). As shown in the enlarged circle and rectangle of Fig. 1B, the shape of this time series is comparable at different zoom levels, but is not exactly the same as for the Koch snowflake of Fig. 1A. The same is true for EMG data, as shown in the enlarged circles of Fig. 2A-B. Both the stock prices and EMG do not satisfy the condition of *strict* self-similarity and should hence better be defined as *statistically* self-similar or, as Mandelbrot suggested, self-affine (Mandelbrot 1985). For this and other reasons, care should be taken when conducting fractal analysis of physiological data, not to misunderstand the sources of variation (Gálvez Coyt et al. 2013; Jelinek et al. 2005; Kesić and Spasić 2016; Müller et al. 2017). For instance, the Higuchi’s fractal dimension (HFD, see methods) is a popular metric for the estimation of fractal dimension in neurophysiology (Kesić and Spasić 2016). Its values, however, can be influenced by numerical perturbations (Liehr and Massopust 2020), noise (Müller et al. 2017), and periodicity of the signal (Gálvez Coyt et al. 2013).

**Fig. 2.**
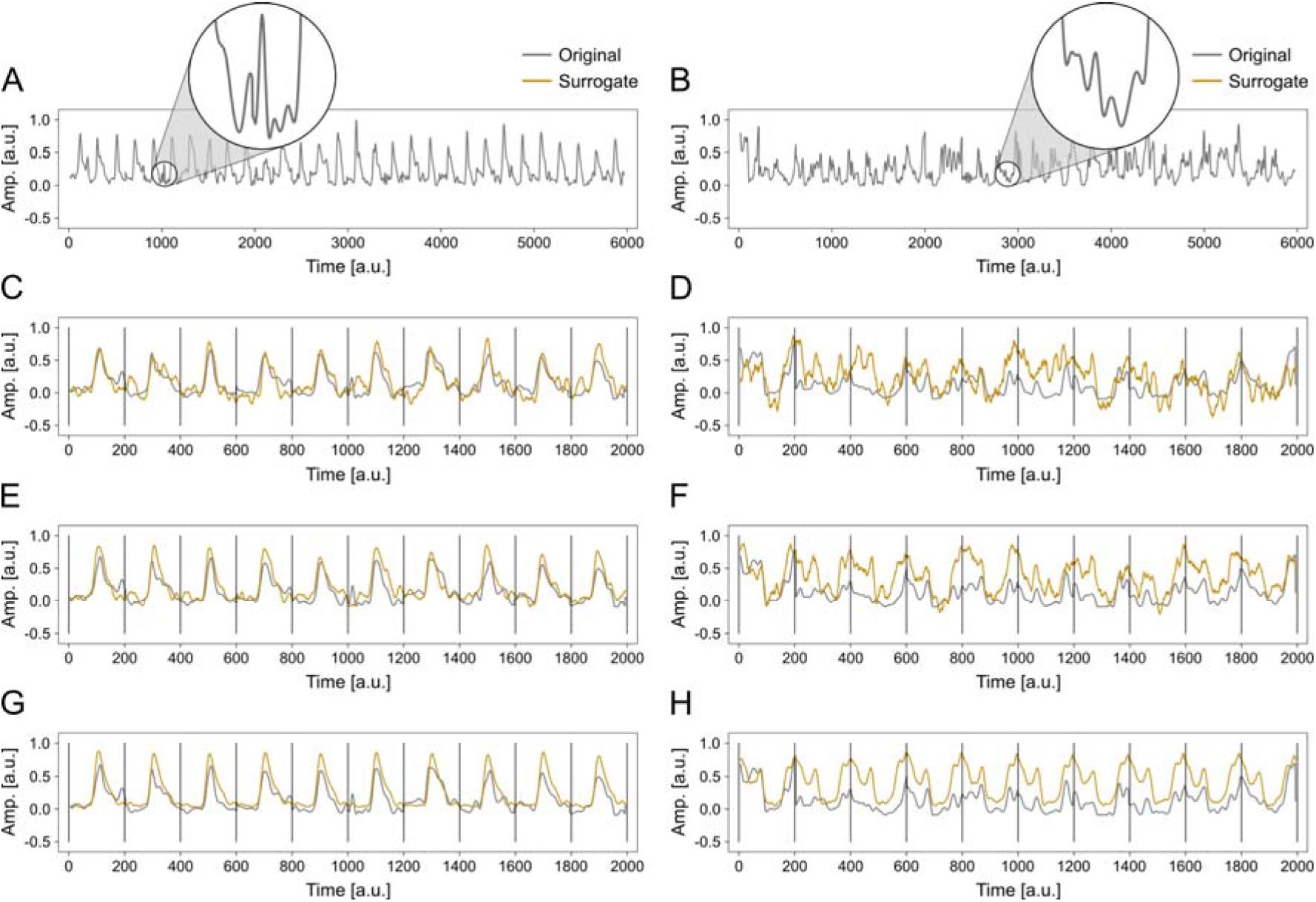
A-B. Two original time series used as baseline and obtained from factorization of electromyographic signals recorded during murine normal (A) and perturbed (B) locomotion. C-D. Detail (first 2000 points) of the original (gray) and one representative surrogate (yellow) data series for normal (C) and perturbed (D) walking with the same mean, variance, autocorrelation and power spectrum of the relevant baseline. E-F. The same as in C-D, but with amplitude of the fluctuations around the mean reduced to 50% of the original. G-H. The same as in C-D, but with amplitude of the fluctuations around the mean reduced to 10% of the original. Time (x-axis) and amplitude (y-axis) of this representative section of a time series are reported in arbitrary units (a.u.).

Here, we propose to use HFD and H as descriptors of changes in EMG-related time series when external perturbations are added to mouse locomotion. We show the sensitivity of the HFD and H to well defined changes caused by targeted manipulation of surrogate data (Dingwell and Cusumano 2000; Theiler et al. 1992). Moreover, we investigated the sensitivity of the HFD and H to physiological changes in motor primitives caused by external perturbations. Finally, we offer some evidence-based suggestions to make the most out of physiological data’s fractal analysis.

## Materials and Methods

### Baseline data

All procedures were performed according to the guidelines of the Canadian Council on Animal Care and approved by the local councils on animal care of Dalhousie University with protocol #19-014. Five adult wild-type mice (age 67 ± 8 days) with C57BL/6 background walked at 0.3 m/s on a custom-built treadmill (workshop of the Zoological Institute, University of Cologne) capable to administer perturbations through quick mediolateral displacements and accelerations of the belt. EMG data was recorded from the hindlimb as previously reported (Akay et al. 2014; Santuz et al. 2019). Basic muscle activation patterns, or motor primitives (Dominici et al. 2011; Santuz et al. 2017), were extracted from EMG using the muscle synergies approach (Bernstein 1967; Bizzi et al. 1991) via non-negative matrix factorization (Lee and Seung 1999; Santuz et al. 2019) using custom R scripts (R v3.6.3, R Core Team, 2020, R Foundation for Statistical Computing, Vienna, Austria). Of the extracted primitives, we arbitrarily chose one for normal and one for perturbed walking, both representative of all five recorded animals. The two time series had the same length (i.e. 6000 data points, see Fig. 2A-B) and described one of the basic muscle activation patterns extracted from 30 gait cycles. Each cycle was time-normalized to 200 points (Santuz et al. 2019): 100 points assigned to the stance (when the limb was in contact with the ground) and 100 to the swing phase (when the limb was swinging in the air).

### Surrogate data

From the two baseline time series, we generated three datasets using a modified version of the surrogate data method (Dingwell and Cusumano 2000; Theiler et al. 1992) using custom R scripts. For each of the two baseline series we: 1) calculated the mean gait cycle; 2) subtracted the mean signal from the baseline to obtain the fluctuations around the mean; 3) computed the fast Fourier transform of the obtained fluctuations; 4) randomized the phase spectrum to generate phase-randomized surrogates of the original fluctuations; 5) computed the real part of the inverse Fourier transform to return the new fluctuations; 6) added the new fluctuations to the mean signal to get the surrogate time series; 7) repeated the previous points 1000 times. This procedure allowed us to obtain two sets of 1000 time series each, one for normal and one for perturbed walking. Any of the 1000 series had the same mean, variance, autocorrelation and power spectrum of the relevant baseline (BSLN, Fig. 2C-D). Additionally, we generated two other data sets by adding a multiplication factor to the real part of the inverse Fourier transform (point 5 of the numerical procedure). We multiplied the fluctuations by 0.5 (BSLN50, Fig. 2E-F) and 0.1 (BSLN10, Fig. 2G-H), operation that produced new time series that fluctuated around the mean with half and one tenth of the original amplitude, respectively. This resulted in more “regular” data sets, in which each gait cycle was more similar to the others as compared to the baseline. In total, we generated 3000 time series of 6000 points each for normal walking (1000 BSLN, 1000 BSLN50 and 1000 BSLN10) and as many for perturbed walking, for a total of 6000 surrogate time series.

### Higuchi’s fractal dimension

The Higuchi’s fractal dimension (HFD) was calculated as previously reported (Santuz et al. 2020). Briefly, using moving windows of different size k, the length L defined by Higuchi (Higuchi 1988) was determined as shown in Fig. 3. L is not an Euclidean length, but is calculated using the y-axis values of the time series at different intervals k: first one takes the sum of the absolute values of their differences; then, the obtained numbers are normalized to the length of the considered sub-series, to make L comparable at different k. This definition implies that L = 0 when taken between two points with the same ordinate (i.e. when L is “horizontal”) and L → ∞ if the two points are very far apart on the y-axis (i.e. when L tends to be “vertical”). Following Higuchi’s definition, if the time series is fractal then the logarithmic average of all obtained lengths is proportional to the logarithm of the window size, with a slope that is the HFD (see Fig. 4A-C). HFD values are independent on the amplitude of the signal and range from 1 (e.g. for a straight line) to 2 (e.g. for a filled rectangle), with HFD = 1.5 for random Gaussian noise; higher HFD indicate more complex data (Anmuth et al. 1994; Higuchi 1988; Kesić and Spasić 2016).

**Fig. 3.**
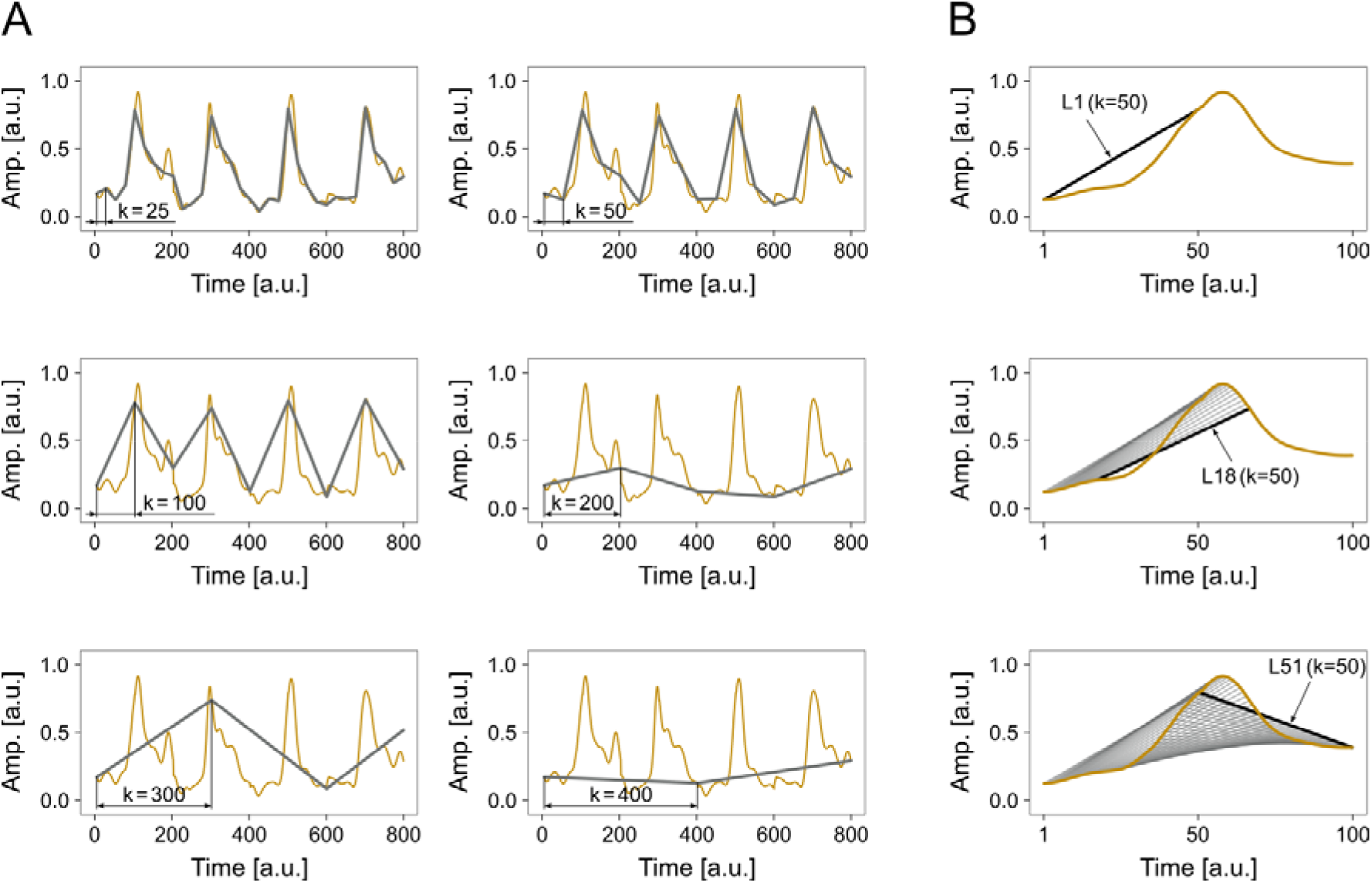
The Higuchi’s fractal dimension is computed by calculating the non-Euclidean Higuchi’s length L (Higuchi 1988) at various time resolutions (panel A) and with different starting points (panel B). Time (x-axis) and amplitude (y-axis) of this representative section of a time series are reported in arbitrary units (a.u.). A. The gray lines represent the original time series sampled with increasingly rough time resolution (i.e. taking one point every 25, 50, 100, 200, 300, or 400 points of the original series). B. The procedure for k = 50 is shown. First, L is computed between point 1 and 50; then between point 2 and 51, between point 3 and 52, etc. The final length for that time resolution k is the average of all the individual lengths. Due to its non-Euclidean definition, L = 0 when horizontal.

**Fig. 4.**
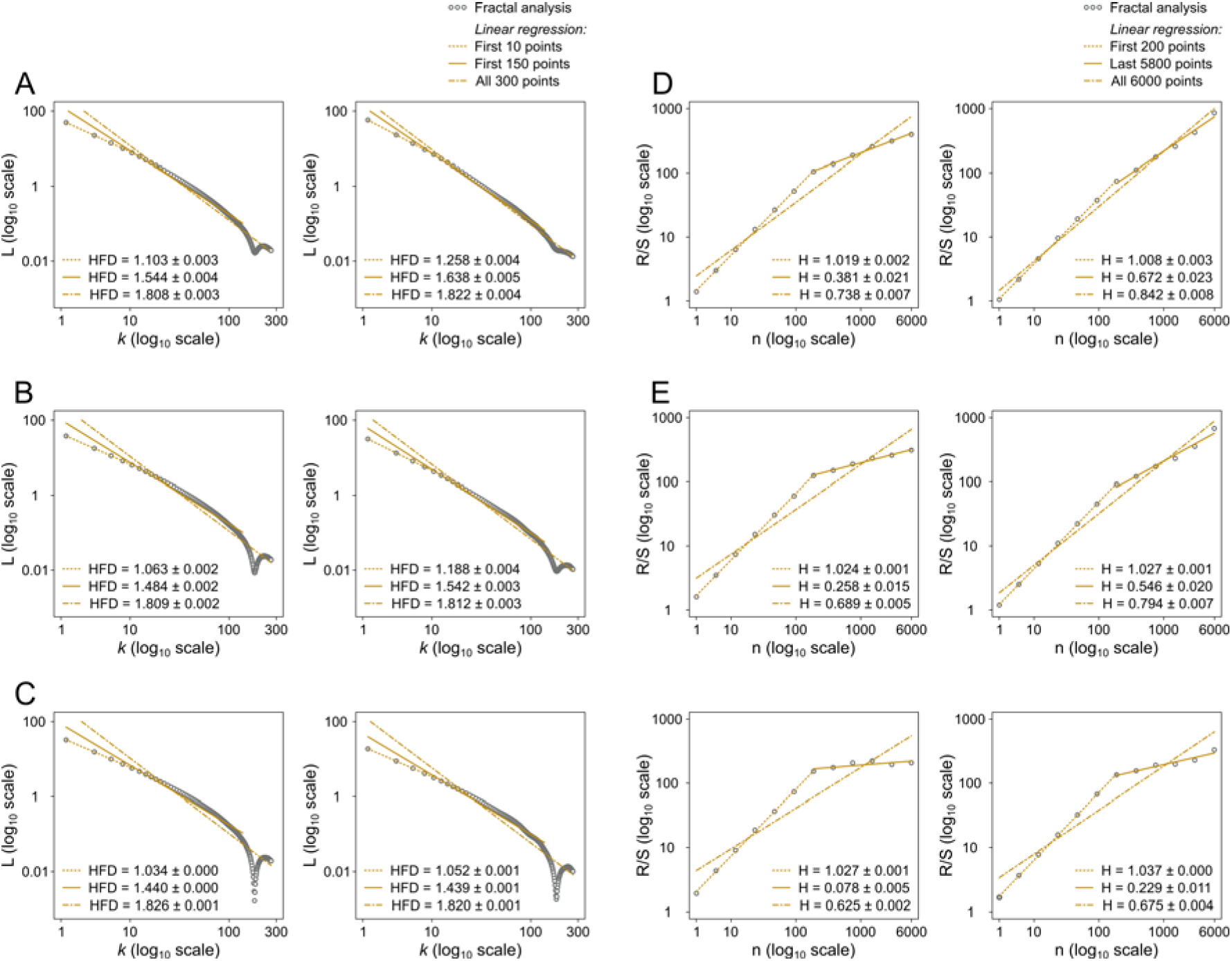
A-C. Higuchi’s log-log plot for 1000 surrogate time series obtained from normal (left) and perturbed (right) locomotion with fluctuations around the mean reduced to 50% (B) and 10% (C) of the original (A). Mean absolute value Higuchi’s fractal dimension (HFD) of the 1000 surrogate data series and standard deviation are reported in each plot after linear regression of the first 10, 150 and all 300 data points. D-F. Hurst’s log-log plot for 1000 surrogate time series obtained from normal (left) and perturbed (right) locomotion with fluctuations around the mean reduced to 50% (E) and 10% (F) of the original (D). Mean Hurst exponent (H) of the 1000 surrogate data series and standard deviation are reported in each plot after linear regression of the first 7, last 6 and all data points.

### Hurst exponent

The Hurst exponent (H) was calculated following the rescaled range (R/S) approach (Mandelbrot and Wallis 1969). We proceeded as follows: 1) calculated the mean of the considered time series of length N; 2) subtracted the sample mean from the original series to normalize the data; 3) calculated the cumulative sum of the obtained series; 4) found the range R of the cumulative sum as the difference between its maximum and minimum values; 5) calculated the standard deviation S; 6) computed the rescaled range R/S; 7) repeated the previous for n = N/2, N/4, N/8… and until a minimum of n=2; 8) calculated H as the slope of the log(n) vs. log(R/S) curve (see Fig. 4D-F). H can vary between 0 and 1. For 0.5 < H < 1, in the long term high values in the series will be probably followed by other high values and a positive or negative trend is visible, similar to when there is a clear increase or decrease of a stock price (see example in Fig. 1B, H_[615, 672]_); in other words, the series is persistent or has long memory (Gneiting and Schlather 2004; Mandelbrot 1983). For 0 < H < 0.5, in the long term high values in the series will be probably followed by low values as in EMG-related data from locomotion, with a frequent switch between high and low values (see example in Fig. 1B, H_[1, 712]_); in other words, the series is anti-persistent (Gneiting and Schlather 2004; Mandelbrot 1983). Examples are given in Fig. 1B. A Hurst exponent of 0.5 indicates a completely random series without any persistence (Mandelbrot 1983; Qian and Rasheed 2004).

### Statistics

To evaluate differences in HFD and H, we estimated the 95% confidence intervals (CI) of the compared sets of surrogate data using the 2.5% sample quantile as the lower bound and the 97.5% sample quantile as the upper bound. Ten thousand resamples with replacement for each parameter were used to estimate the CI (Santuz et al. 2019). Moreover, we calculated the effect size Hedges’ g. Differences were considered significant when the zero was lying outside each CI. The level of significance was set at α = 0.05. All statistical analyses were conducted using custom R scripts.

### Data availability

In the supplementary data set accessible at Zenodo (DOI: 10.5281/zenodo.3764734) we made available: a) the metadata file “metadata.dat”; b) the baseline signals file “baseline_data.RData”; c) the R script “fractal_analysis.R” to calculate the H and HFD of the baseline data and produce the log-log plots. Explanatory comments are profusely present throughout the script and in the metadata file.

## Results and Discussion

We found that both HFD and H values of almost periodic physiological time series depend on several variables. Care should be taken when conducting this kind of fractal analysis and in the next paragraphs we will present and discuss our findings, concluding with some suggestions.

### Rationale

Results are summarized in Fig. 4. The top panels (Fig. 4A for normal and Fig. 4D for perturbed walking) describe the HFD and H calculations at different window lengths for surrogate data with the same fluctuations around the mean as the baseline’s, only randomized in order. The mid panels (Fig. 4B for normal and Fig. 4E for perturbed walking) describe the HFD and H calculations at different window lengths for surrogate data with fluctuations around the mean reduced to half of the baseline’s. The bottom panels (Fig. 4C for normal and Fig. 4F for perturbed walking) show the results after reduction of the fluctuations to just the 10% of the baseline’s. This allowed us to analyze three levels of “regularity” in our surrogate time series: from the lowest (i.e. the original) to the highest (i.e. 10% of the original fluctuations). This concept is visible in Fig. 2.

### Quasi-periodic time series

Locomotion dynamics are quasi-periodic (Scafetta et al. 2009). This means that when we record some physiological parameters such as muscle activity, their behavior in time will not be as regular as that of a sinusoidal function, but it will likely be close enough to allow the identification of similar cycles (e.g. the steps or the strides). The time series considered in this study have been time-normalized to 200 points assigned to each gait cycle. Thus, their period is 200. As it is visible in Fig. 2, the quasi-periodicity becomes more evident when shifting the attention from BSLN (Fig. 2C-D) to BSLN50 (Fig. 2E-F) and, ultimately, BSLN10 (Fig. 2G-H).

### Higuchi’s fractal dimension

The HFD values (Fig. 4A-C) were significantly different between normal (left) and perturbed (right) walking at all window lengths with absolute values of the Hedges’ g effect size between 3.2 and 96.3. This indicates the ability of HFD to be sensitive to the presence of perturbations during locomotion. However, HFD values increased as the length of the window used for regression increased. This behavior shows that the curve in the log-log plot is not linear as in the case of strictly self-similar data series (Higuchi 1988). Due to the lack of consensus in the literature on how to choose the maximum window length (what is often called k_max_) and select the most linear part of the log-log curve (Kesić and Spasić 2016), extreme care should be taken when calculating the HFD of quasi-periodic time series. It is not surprising that some authors suggest to only consider the first two points in the log-log plot, so as to make sure that the selected portion is trivially linear (Gneiting et al. 2012). Moreover, a discontinuity appeared in the neighborhood of the 200^th^ time point, bigger in normal as compared to perturbed walking and increasing from BSLN to BSLN10. This discontinuity is responsible for a sharp increase in the slope (which is the HFD) of the regression lines that include it, with an effect that increases with the “regularity” of the time series. This is due to the definition of the Higuchi’s length (Higuchi 1988), which tends to zero when taken between two points that have close ordinate component or, in other words, when it is almost horizontal in the example of Fig. 3B (Gálvez Coyt et al. 2013; Liehr and Massopust 2020).

### Hurst exponent

The H values (Fig. 4D-F) were significantly different between normal and perturbed walking at all window lengths with absolute values of the effect size between 5.1 and 45.6. This indicates the ability of H to be sensitive to the presence of perturbations during locomotion. However, H values (or the slope of the considered curve in the log-log plot) were higher for minimum window lengths smaller than 200 points and lower for minimum window lengths bigger than 200 points, divided by what is called a crossover point (Mandelbrot 1985). This not only shows the susceptibility of H to the chosen window length, but also that H is influenced by the presence or absence of quasi-periodic elements in the time series. This can be explained by comparing the properties of H and HFD. For self-similar series, the fractal dimension is 2-H (Mandelbrot 1985), thus lower H (or higher HFD, as stated above and in the methods) indicate more irregular or complex data (Higuchi 1988; Kesić and Spasić 2016). For self-affine series such as those treated in this study, fractal dimension and H are not necessarily linearly correlated and can exhibit different local and global values (Gneiting and Schlather 2004; Mandelbrot 1997). In the words of Gneiting and Schlather (Gneiting and Schlather 2004): “*[…] fractal dimension and Hurst coefficient are independent of each other: fractal dimension is a local property, and long-memory dependence is a global characteristic*.”. In fact, if the data shows any short-range dependence as in this case, it is commonly suggested to exclude the low end of the plot for the estimation of H (Taqqu et al. 1995).

### Concluding remarks

When dealing with physiological data, it is tempting to use fractal analysis for estimating the regularity, complexity or persistence of time series (Kesić and Spasić 2016; Losa et al. 2005). Here we showed that the values of two common metrics for fractal analysis, HFD and H, are exceptionally sensitive to changing in EMG-derived data when external perturbations are applied to murine locomotion. However, HFD and H also strongly depend on the parameters selected for the calculations and the presence or absence of quasi-periodic elements in the time series. In order to work around those issues, we suggest to: a) calculate HFD by using only the most linear part of the log-log plot, which in our case correspond to maximum the first 10 data points, and b) calculate H by using a smallest window bigger than the period given by the length of the individual gait cycles.

## Acknowledgments

t.b.d.

## Author contributions

Conceptualization: A.S., and T.A.; Data curation: A.S.; Formal analysis: A.S.; Investigation: A.S.; Methodology: A.S., and T.A.; Project administration: A.S., and T.A.; Resources: A.S., and T.A.; Software: A.S.; Supervision: T.A.; Validation: A.S., and T.A.; Visualization: A.S.; Writing – original draft: A.S., and T.A.; Writing – review & editing: A.S., and T.A.

